# scCapsNet-mask: an automatic version of scCapsNet

**DOI:** 10.1101/2020.11.02.365346

**Authors:** Lifei Wang, Jiang Zhang, Jun Cai

## Abstract

**Summary:** Recently we developed scCapsNet, an interpretable deep learning cell type classifier for single cell RNA sequencing data, based on capsule network. However, the running process of scCapsNet is not fully automatic, in which a manual intervention is required for getting the final results. Here we present scCapsNet-mask, an updated version of scCapsNet that utilizes a mask to fully automate the running process of scCapsNet. scCapsNet-mask could constrain the internal parameter coupling coefficients and result in a one to one correspondence between the primary capsule and type capsule. Based on those bijective mapping between primary capsule and type capsule, the model could automatically extract the cell type related genes according to weight matrix connecting input and primary capsule, without a need for manual inspection of the relationship between primary capsules and type capsules. The scCapsNet-mask is evaluated on two single cell RNA sequence datasets. The results show that scCapsNet-mask not only retains the merits of the original scCapsNet with high classification accuracy and high interpretability, but also has the virtue of automatic processing.

## Introduction

Single Cell RNA sequencing (scRNA-seq) could measure gene expression levels in individual cells(Chen, et al., 2019). Recent advances in scRNA-seq require more computational analysis methods. Deep learning model is a proper tool to deal with vast and complex data such as RNA-seq data (Eraslan, et al., 2019; Lopez, et al., 2018), but lacks of interpretability (Almas Jabeen, 2017). Therefore, there is a need for a model that utilizes the deep learning method with increasing interpretability in order to reveal the biological meaning hiding behind the model (Ding, et al., 2018; Hung-I Harry Chen, 2018; Lin, et al., 2017).

The CapsNet is a novel deep learning model which is used in the task of digit recognition (Sabour, et al., 2017) and hold the great potential to apply in network biology and disease biology with data from multi-omics datasets (Camacho, et al., 2018).

Previously, we proposed the single cell capsule network (scCapsNet) model, a highly interpretable classifier for dealing with scRNA-seq data adopted from capsule network(Wang, et al., 2020). Through model internal parameters, scCapsNet can not only classify cell subpopulations with high accuracy, but also reveal the related genes of cell subpopulations that determine the process of classification. The coupling coefficient, one of the internal parameters, reveals the relationship between primary capsule and type capsule, and is a matrix with row representing type capsule and column representing primary capsule. Due to the variability of the coupling coefficient, which changes with each round of training, the relationship between primary capsule and type capsule is not fixed and needs manual inspection for downstream process. In order to automate the running process of the scCapsNet, we set the number of primary capsule as the number of type capsule, and then constrain the coupling coefficient by adding a mask in the dynamic routing process of capsule network, inspired by Sparse Transformers(Child, et al., 2019). The constrained coupling coefficient is a diagonal squared matrix and the one to one correspondence between primary capsule and type capsule is achieved. We call this version of the scCapsNet model “scCapsNet-mask”. We apply scCapsNet-mask to scRNA-seq data of mouse retinal bipolar cells (RBC) (Lopez, et al., 2018; Shekhar, et al., 2016) and human peripheral blood mononuclear cells (PBMC) (Lopez, et al., 2018; Zheng, et al., 2017). The results show that the scCapsNet-mask not only retains the merits of the original scCapsNet with high classification accuracy and high interpretability, but also has the virtue of automatic processing.

## Methods

### scCapsNet-mask

The scCapsNet model is adapted from previous CapsNet model. The model contains two parts, one of which is feature extraction part and the other is capsule network part. The feature extraction part consists of several parallel fully connected neural networks using Rectified Linear Unit (ReLU) activation function as the feature extractor. The capsule network part directly used a Keras implementation of CapsNet and the number of the primary capsules is a hyper-parameter of the model. In the scCapsNet-mask, the number of the primary capsules is the same as that of the type capsules, which is the number of cell types in training set. So the coupling coefficient is a square matrix. Then we add a mask to the square matrix of the coupling coefficient, in order to create a one to one correspondence between primary capsule and type capsule. The mask is a diagonal matrix, with on-diagonal elements all being one and the off-diagonal elements all being zero. In each round of the “dynamic routing”, the coupling coefficients would multiply with this mask matrix first, in order to concentrate the weights in the on-diagonal elements and ignore the off-diagonal elements of the square matrix of the coupling coefficient (Fig 1).

**Figure 1.**
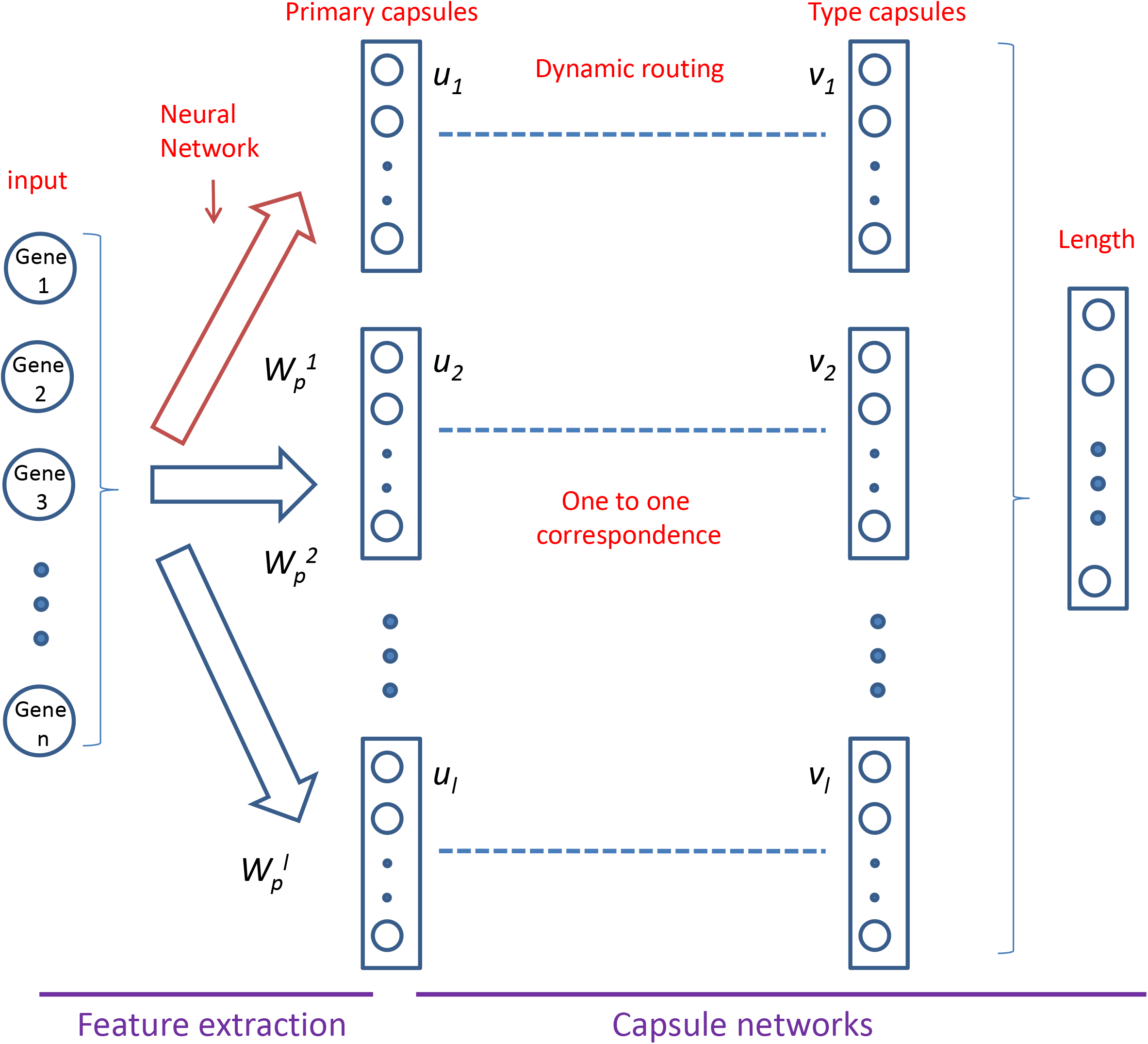
Two-layer architecture of scCapsNet-mask. The first layer consists of *l* parallel fully-connected neural networks for feature extraction from the inputs of single-cell gene expression. The *l* is equal to the number of cell types. The primary capsule of vector u_i_ is the output of the neural network *i*. The subsequent layer is a Keras implementation of capsule networks for classification with a mask added on the parameters of coupling coefficients. The result of the mask is the one-to-one correspondence between primary capsules and type capsules. The length of each type capsule v_j_ represents the probability of single cell x belonging to the corresponding cell type.

## Results

To test our model, two scRNA-seq datasets are used. One consists of mouse retinal bipolar cells (mRBC) profiled by the Drop-Seq technology and the other consists of human peripheral blood mononuclear cells (hPBMC) generated by 10X Genomics. Both datasets are randomly divided into training set and validation set with a ratio of 9:1. The model is run several times to access the prediction accuracy. Similar as the scCapsNet, the validation accuracy of scCapsNet-mask model for mRBC and hPBMC are very high, around 98% and 96%, respectively.

After training, we re-run the model and calculate the average coupling coefficients for each cell type. From the heatmap of the average coupling coefficients of each cell types, we could clearly see that there is one and only one most active element in each average coupling coefficient (Fig 2A, Fig 4). Furthermore, the overall heatmap of the average coupling coefficients for all cell types demonstrate that only the on-diagonal elements are active, and each primary capsule is only related with one type capsule and vice versa (Fig 2B, Fig 4 lower right). These results undoubtedly justify our proposal of effect of the mask, that is, the mask would constrain the weight distribution of the coupling coefficients and concentrate the weights on the on-diagonal elements. After applying the mask, there is a one to one correspondence between primary capsule and type capsule. So the model could automatically find the relationship between the primary capsule and type capsule, without needing manual inspection to indicate which primary capsule responds to the recognition of which cell types.

**Figure 2.**
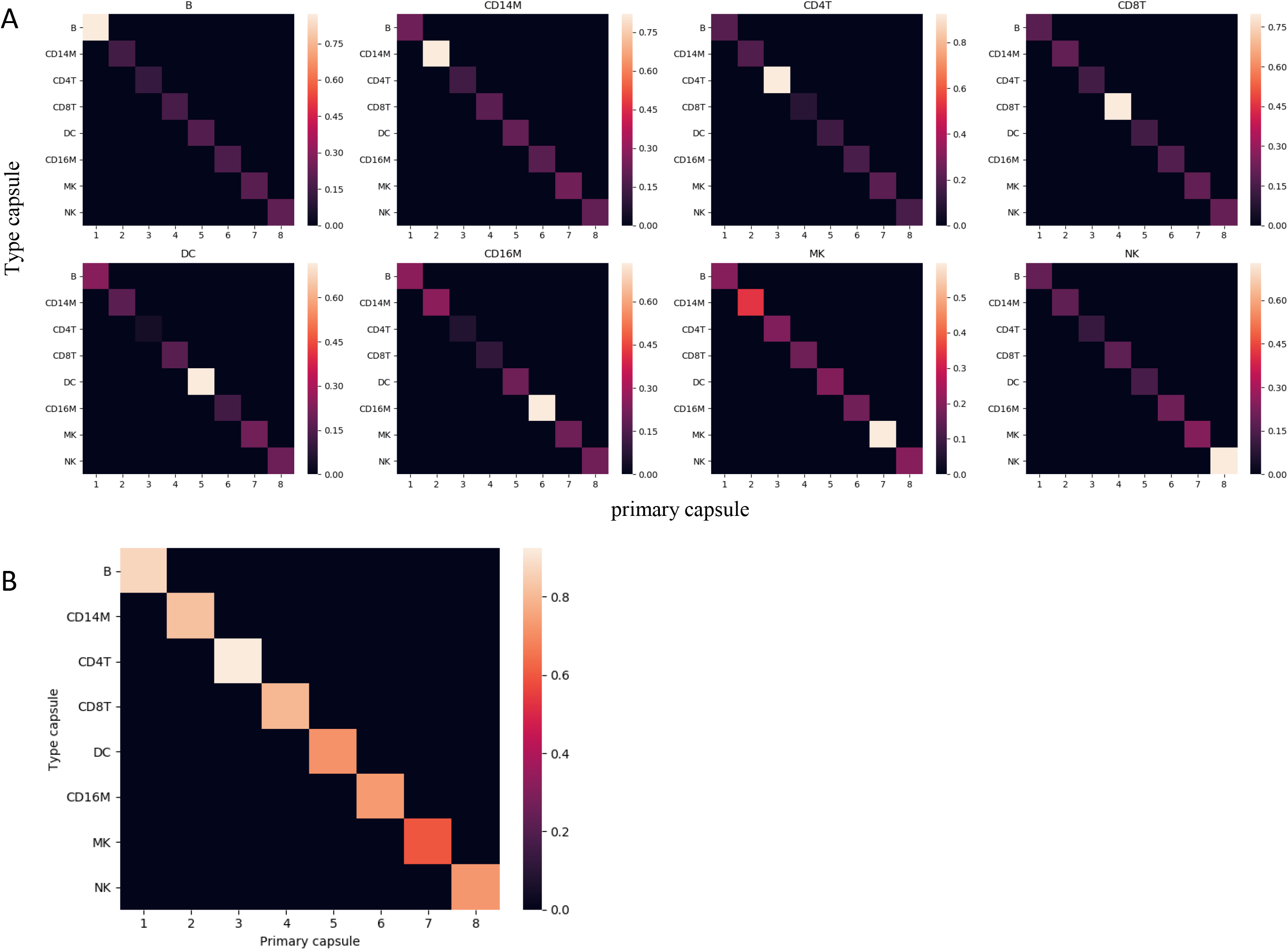
A: The heatmaps of the matrices of averaged coupling coefficients for hPBMC dataset with cell types listed above. The ‘B’, ‘CD14M’, ‘CD4T’, ‘CD8T’, ‘DC’, ‘CD16M’, ‘MK’, and ‘NK’ are abbeveation for B cells, CD14+ monocytes, CD4+ T cells, CD8+ Tcells, dendritic cells, FCGR3A+ monocytes, megakaryocytes, and natural killer cells, respectively. For each heatmap, the row represents type capsules and the column represents primary capsules. B: The overall heatmap of the combining matrix of average coupling coefficient. The combining matrix contains the effective type capsule row in Figure 2A where its recognition type is in accordance with the type of input single cells.

As described for the previous version of scCapsNet, the internal weight matrix in the networks connecting the inputs with each primary capsule would further connect genes from input to each cell type. Each column vector of the internal weight matrix represents a low dimensional representation of each gene. As previously, we perform PCA on column vectors of the internal weight matrix from specific primary capsule (which also mean a specific cell type), and then choose genes according to their coordinates on PC1 (Fig 3B, Fig 6). When we exclude this group of genes in the inputs of the scCapsNet model, a new set of cell-type recognition accuracies is obtained. As we slide the cutoff values along the principal component scores, a set of cell-type recognition accuracy curves is generated. In the accuracy curves plot, the recognition accuracy of one cell type reduces much more sharply than others with the ascending or descending sliding cutoff values (Fig 3A, Fig 5). There exists an appropriate threshold of principal component scores (Fig 3A, Fig 5 dotted line). When we exclude those genes in the inputs of the scCapsNet model, the model almost could not recognize that specific cell type, while the ability to identify other cell types is partially retained or even left intact. These sets of genes, which are selected by model for one specific cell type, contain several well studied bio-markers corresponding to this cell type (Fig 3B).

**Figure 3.**
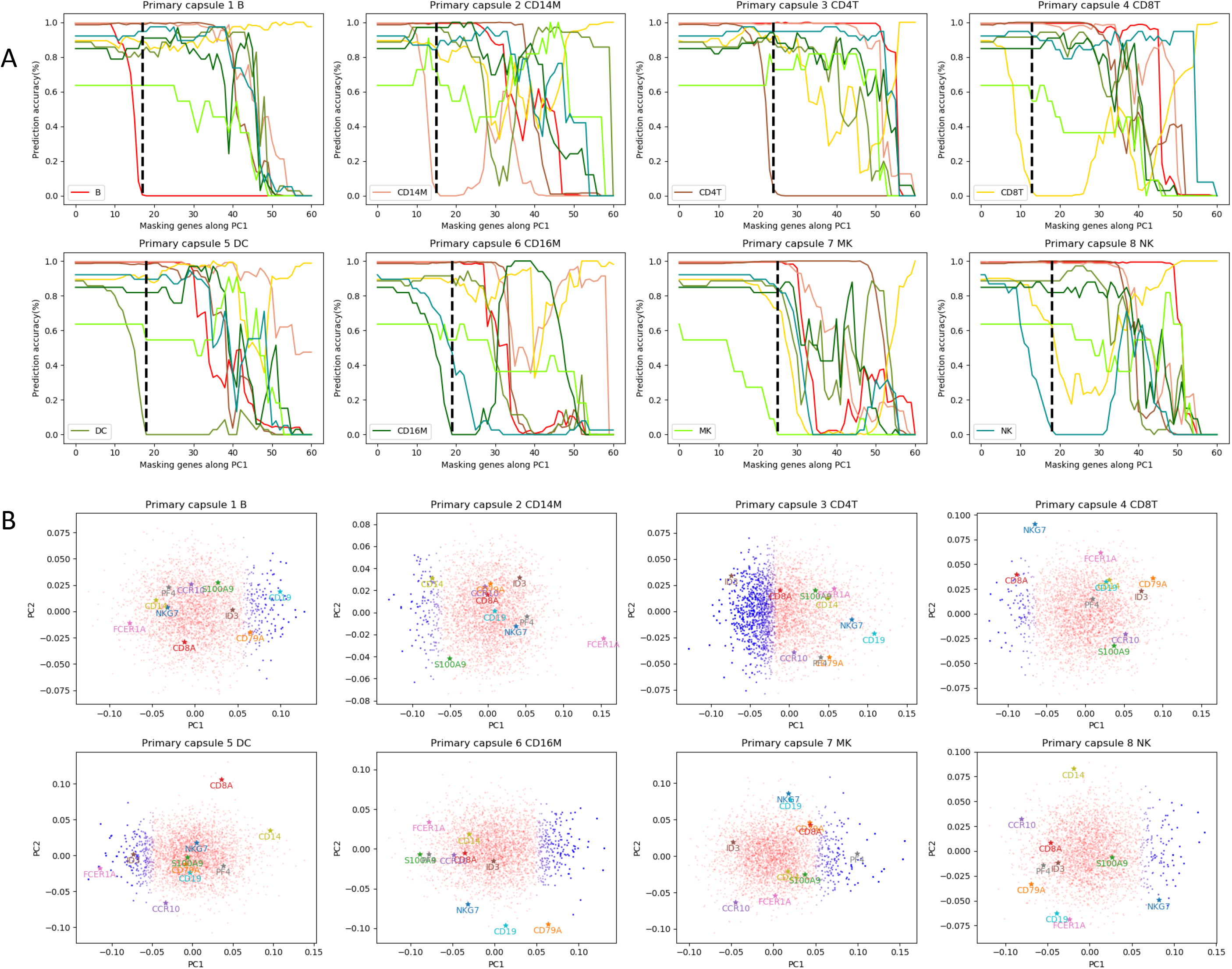
A: The colored changing curves of cell-type recognition accuracies with the ranking genes defined by a sliding cutoff value on the principal component score being excluded in the inputs of the scCapsNet-mask model. The accuracy curve for each cell type is represented by a distinct color. The dotted line defines a group of cell type related genes responsible for cell identification, where the recognition accuracy degrades close to 0 for corresponding cell type listed above but not significantly decreases for any other cell types in general. B: The plot depicts the two-dimensional principal component analysis (PCA) on the weight matrix for the primary capsule. Each dot represents a gene with a rank according to the score of principal components. A group of cell type related genes marked as blue color is defined. The colored stars with labels represent the bio-markers.

**Figure 4.**
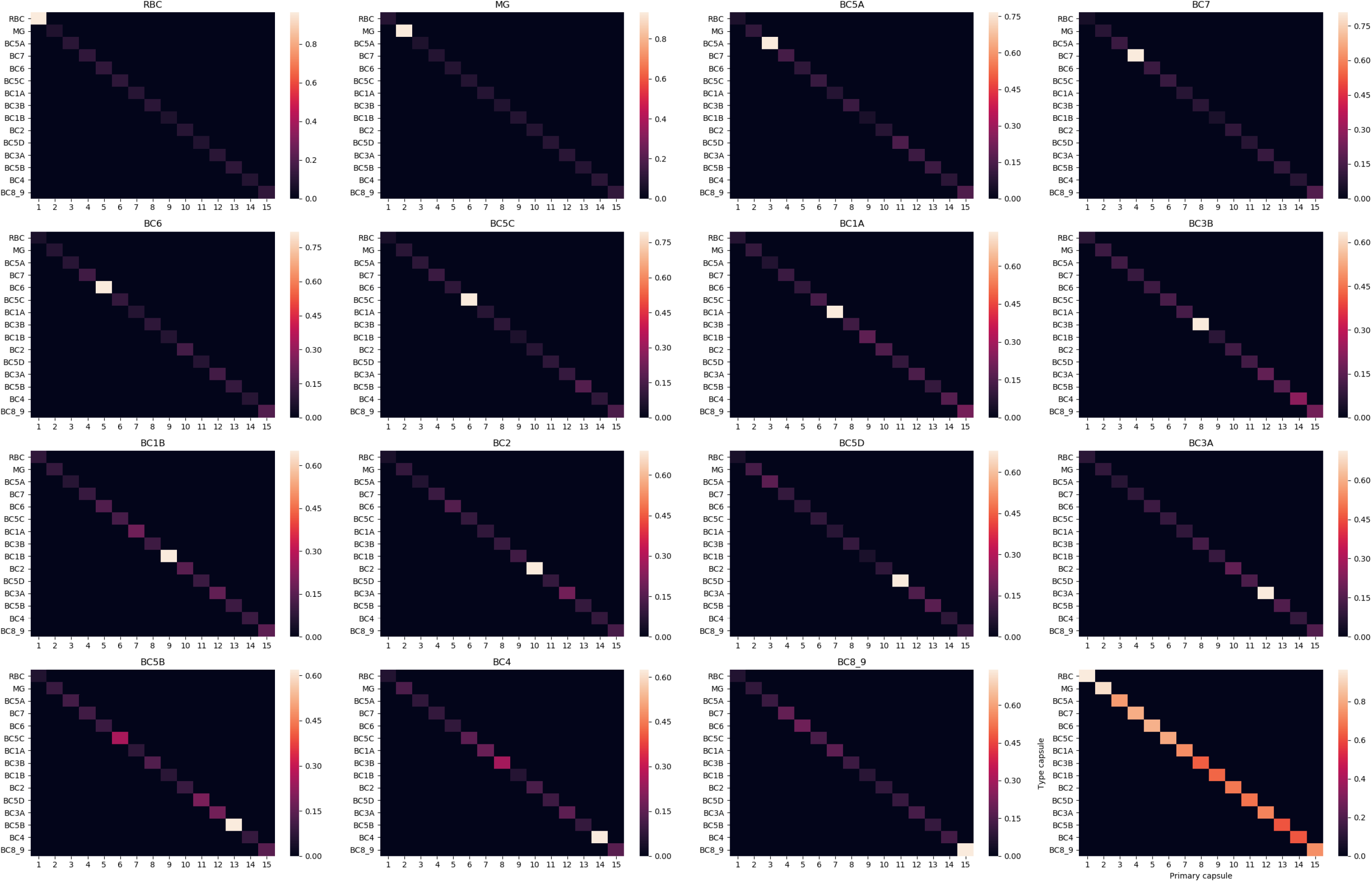
The heatmaps of the matrices of averaged coupling coefficients for mRBC dataset with cell types listed above. For each heatmap, the row represents type capsules and the column represents primary capsules. The last heatmap represents the overall heatmap of the combining matrix of average coupling coefficient. The combining matrix contains the effective type capsule row in previous 15 heatmap where its recognition type is in accordance with the type of input single cells.

**Figure 5.**
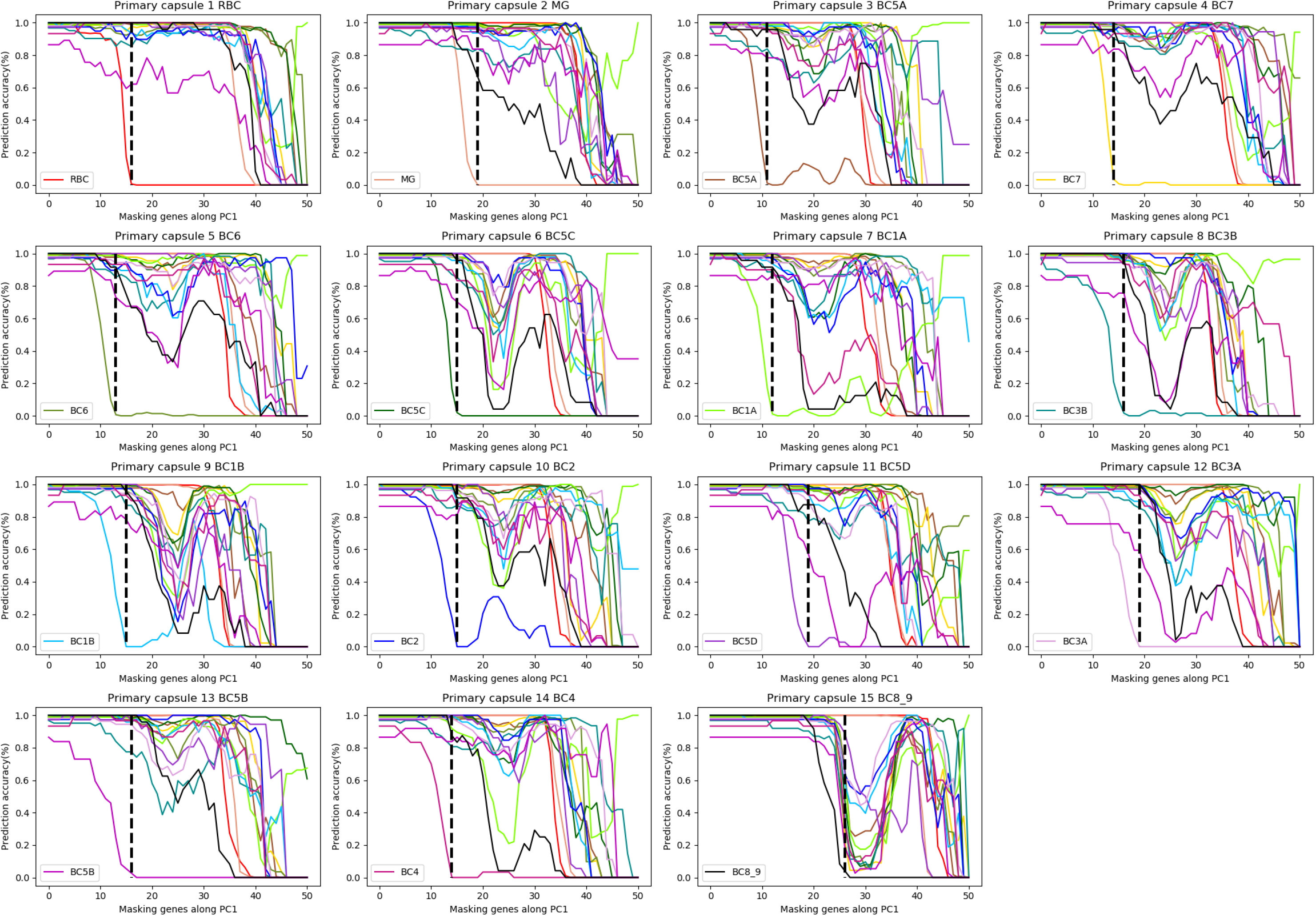
The colored changing curves of cell-type recognition accuracies with the ranking genes defined by a sliding cutoff value on the principal componentscore being excluded in the inputs of the scCapsNett-mask model. The accuracy curve for each cell type is represented in a distinct color. The dottedline defines a group of cell type related genes responsible for cell identification, where the recognition accuracy degrades close to 0 for corresponding cell type listed above but not significantly decreases for any other cell types in general.

**Figure 6.**
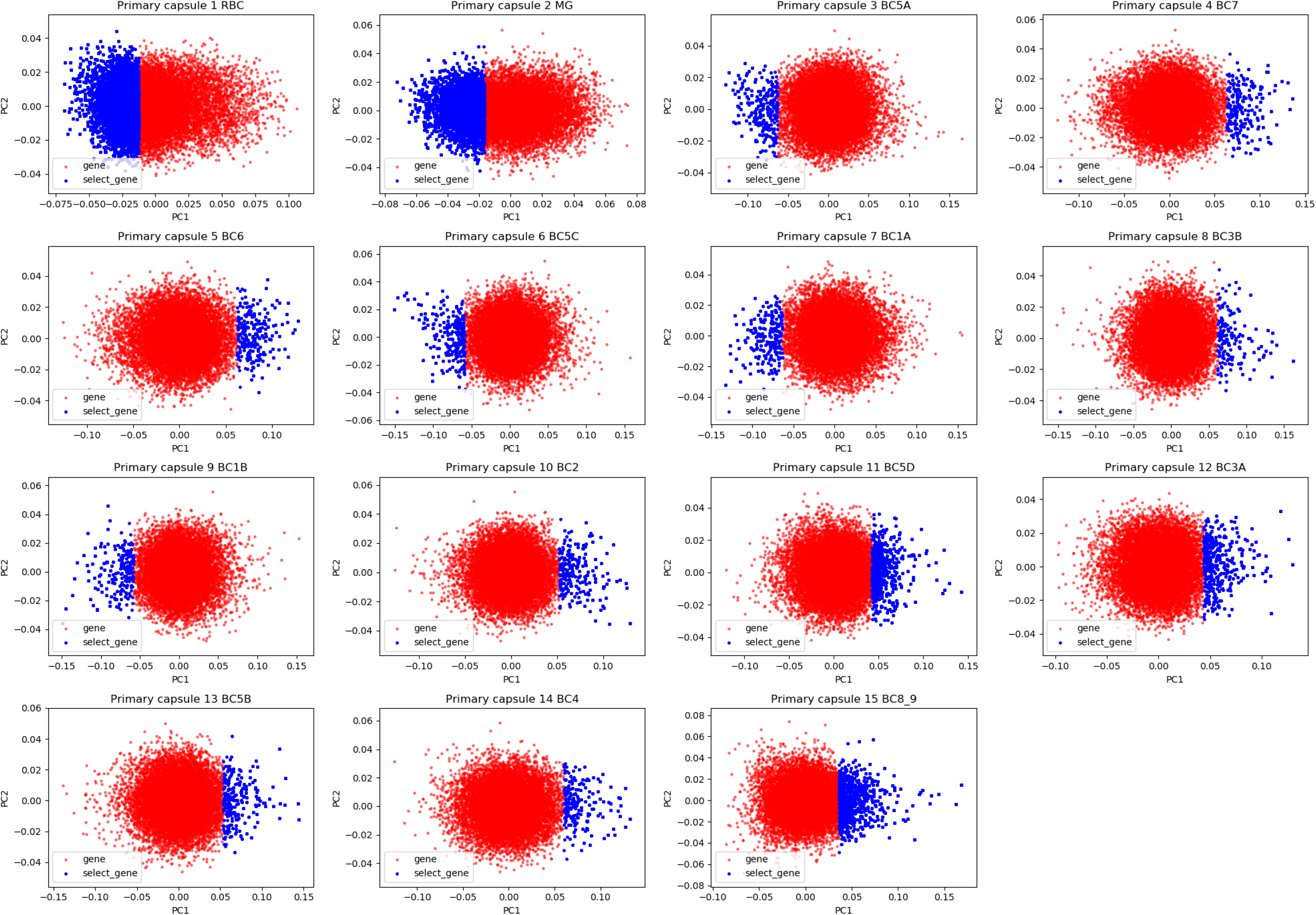
The plot depicts the two-dimensional principal component analysis (PCA) on the weight matrix for the primary capsule. Each dot represents a gene with a rank according to the score of principal components. A group of cell type related genes marked as blue color is defined.

Overall, scCapsNet-mask constrains the coupling coefficients, resulting in a bijective mapping between primary capsule and type capsule. As original scCapsNet, the PCA on column vectors of the internal weight matrix from specific primary capsule could reveal several cell type related genes, rendering scCapsNet-mask automatic and interpretable.

